# Vertical sleeve gastrectomy in rodents: impact on metabolic, cardiovascular and hepatic functions of mothers and perinatal consequences

**DOI:** 10.1101/2024.11.12.623181

**Authors:** Cyrielle Payen, Abigaëlle Guillot, Julien Chaigneau, Lucile Bichot, Eugénie Testa, Jennifer Bourreau, Géraldine Gascoin, Emilie Vessières, Carolyne Proux, Valérie Moal, Laurent Loufrani, Françoise Schmitt, Céline Fassot

## Abstract

**Background:** Maternal obesity is a well-known risk factor for obstetrical complications during pregnancy, and cardiovascular and metabolic pathologies in the offspring. In the last two decades, bariatric surgery has emerged as the most effective treatment for severe obesity, and is known to improve fertility, leading to a rise in procedures among women of childbearing age.

**Objectives:** This study investigates the impact of pre-conceptional sleeve gastrectomy (SG) on metabolic, cardiovascular and hepatic functions of mothers at short and middle terms after surgery, and its perinatal consequences.

**Methods:** Sprague-Dawley female control rats (CM) received a standard diet, while others were fed a high-fat, high-sugar diet (HFHS). HFHS rats were divided into obese mothers (OM) and sleeve gastrectomy mothers (SM). One month after surgery, all groups underwent gestation and subsequent lactation Metabolic function was assessed at 1 and 4 months after surgery; cardiovascular and hepatic functions were evaluated only 4 months after SG.

**Results:** From 1 month post-surgery, SM showed normalized leptin and insulin levels compared to OM. Moreover, SG normalized body weight associated with a decreased white adipose tissue weight and adipocyte size at 4 months post-surgery. Interestingly, although steatosis has been detected in HFHS rats, SM exhibited only hepatic fibrosis at 4 months post-surgery. Moreover, SM showed a hypertrophic remodeling of thoracic aorta without effect on its vasoreactivity but along with endothelial dysfunction of mesenteric artery, associated with a stronger involvement of COX/PGI2 pathway. In the offspring, obesity in mothers was associated to a decreased liter size, reversed by sleeve gastrectomy without any effect on the birth weight of pups.

**Conclusion:** While this study reports an improvement of metabolic function at short- and medium-terms after bariatric surgery, our results highlight an alteration of hepatic and vascular functions by sleeve gastrectomy, with minimal effect on the offspring in the perinatal period.

## Introduction

According to the World Health Organization, more than 650 million adults were classified as people with obesity in 2016, 15% of whom are women [1]. The management of obesity and its associated comorbidities (dyslipidemia, diabetes mellitus, metabolic dysfunction-associated fatty liver disease-MAFLD, hypertension) is considered a major public health issue.

Knowing that up to 50% of women of reproductive age and 20-25% of pregnant women in Europe present overweight or obesity [2], obesity in women of child-bearing age is of particular concern. Indeed, the DOHaD concept (Developmental Origins of Health and Disease) makes the periconceptional and intrauterine periods a key window during which nutritional stress will have a lasting impact on fetal development and future health [3]. Maternal obesity is associated with considerable maternal perturbations such as reduced fertility and obstetric complications (preeclampsia, gestational diabetes mellitus), leading to detrimental short-term outcomes for both mother and neonate [4]. Furthermore, maternal obesity impacts fetal development (preterm birth, macrosomia, early neonatal death), increasing the risk for the child to develop metabolic and cardiovascular disorders during adulthood [5–7].

In women with severe obesity, physical activity and / or dietetic interventions are not enough to significantly and durably reduce body weight and then improve postnatal health in children [8]. For the past 20 years, bariatric surgery has emerged and is currently considered as the most effective treatment for severe obesity in order to improve women fertility and maternal / fetal comorbidities [9]. Thus, the incidence of bariatric surgery has increased among obese women of child-bearing age before pregnancy [10]. Different surgical procedures have been developed to reduce body weight. The most widespread procedures are a restrictive and malabsorptive surgery with the Roux-en-Y gastric bypass (RYGB), considered as the gold standard of bariatric surgery [11,12] and a more recent procedure, the vertical sleeve gastrectomy (SG) [13], which has currently become the most practiced. Even if RYGB is more efficient than the other procedures in the midterm [14], the technical simplicity of SG associated to its lower incidence on surgical complications ensures its attractiveness [15].

Current evidences indicate that pre-conceptional bariatric surgery improves maternal and fetal prognosis [16]. Indeed, recent studies highlighted the reduction of several obesity-related pregnancy complications (perinatal outcomes, gestational diabetes mellitus, preeclampsia, macrosomia) after bariatric surgery [17], but at the cost of a strict nutritional monitoring of the mother [18]. However, it appears that children born to operated mothers are at increased risk of prematurity or intrauterine growth retardation [17].

To study the impact of sleeve gastrectomy on mothers and on offspring’s perinatal outcomes, we developed an experimental model on rats. We evaluated metabolic and vascular functions of the mothers at short and middle terms after surgery and perinatal issues of the different groups of pups.

## Methods

### Animal model

Experiments were conducted in accordance with the institutional guidelines (3R principles) and the recommendations for the care and use of laboratory animals of the French Ministry of National Education, Research and Innovation. The protocol received the agreement APAFIS#10697-2017091422557044 v1.

35 Female Sprague–Dawley rats, ∼ 6 weeks of age, were purchased from Janvier Labs (Le Genest S^te^ Isle, France). The animals were housed in individual cages (1 500 cm3 for 3 animals 400-600g) within a controlled temperature room (21-23°C) and maintained under a 12-hr light/dark cycle with free access to water and food. After 7 days of acclimation in the laboratory animal husbandry unit (SCAHU, Angers, France), 11 female rats were randomly assigned to the normal-weight control group (*Control*) and were fed *ad libitum* a standard show diet (8.4% fat, 19.3% protein, 72.4% carbohydrate [w/w], 13.97kJ g-1, diet code: 3430, Serlab, Compiègne, France); the other female rats were fed a high-fat diet (22.6% fat, 23.0% protein, 35% carbohydrate [w/w], 16.73 kJ g-1, diet code: 824053, SDS, Dietex International Ltd., Witham, UK) associated with sweetened condensed milk (55% simple sugar, 8% fat, 8% protein, Nestlé, France) during minimum 8 weeks (*HFHS* group). Obesity was assessed by weight gain and metabolic disorders (calories intake, plasma analysis, hepatic structure; supplemental data, Figure 1S). *HFHS* female rats were then randomly assigned for surgery in one of the 2 following groups: obese mothers (*Obese*) or obese mothers with vertical sleeve gastrectomy (*Sleeve*). After surgery, each female rat was fed with its assigned food until the end of the follow-up.

### Surgical procedures

Surgical interventions were performed after 4 hours fasting, under isoflurane general anesthesia (3.5% v/v in 20% O2 and 80% air). Immediate pre-operative care included subcutaneous injection of 1.5 ml serum saline, 0.1 mg/Kg buprenorphine, 0.5 mg/Kg metoclopramide, 30 mg/Kg amoxicillin/clavulanic acid and 0.175 mg/Kg Fercobsang®.

All the animals of the 3 groups were subjected to a longitudinal median laparotomy, and to a liver biopsy of the median lobe. The abdominal wall of *Control* and *Obese* animals was then closed in two layers using a continuous absorbable suture (V-Loc™ 3/0, Covidien/Medtronic, Paris, France). For the group undergoing sleeve gastrectomy: after the liver biopsy, the right gastric artery was ligated at the bottom of the great curvature of the stomach, and a resection of about 70% of the gastrich pouch including the rumen in totality was performed. The remaining part of the stomach was then tubularized and closed using two subsequent continuous sutures of 6/0 Prolene (Peters Surgical, Bobigny, France). Finally, the abdominal cavity was closed as described previously for the *Control* group.

During the three first postoperative days, they received subcutaneous hydration (1.5 ml serum saline), analgesia (buprenorphine 0.1 mg/Kg), antibioprophylaxis (30 mg/Kg amoxicillin/clavulanic acid), iron supplementation once a day and were fed with GelDiet Energy (Safe, Augy, France). Then all animals were given water supplemented with multivitamins (Vita Rongeur, Virbac France, Carros, France) until the end of the protocol.

One month after recovery, the efficiency of bariatric surgery was assessed by body weight evolution, calories intake, insulin sensitivity and plasma analysis (glycemia, insulin, leptin, triglycerides, cholesterol). Then, each female was mated with a non-obese Sprague-Dawley male to analyze the impact of obesity +/- sleeve gastrectomy on litters (sex-ratio, birth weight and weight gain before weaning) of offspring from control mothers (CMO), obese mothers (OMO) and sleeve mothers (SMO). The experimental design of the study was summarized in supplemental data (Fig. 2S).

Four months after surgery, insulin sensitivity and glucose tolerance were evaluated, as 24h-calories intake. Mothers were then euthanized by carbon dioxide inhalation. For each animal, thoracic aorta and third-order mesenteric arteries free of fat and adhering connective tissues were gently dissected and placed in a physiological salt solution for vascular reactivity study and histomorphometric study. Blood samples, white adipose tissues and liver were also collected.

### 24h-calories intake

Before surgery and 1 month after it, female rats were placed for 24h in individual metabolic cages (Techniplast France, Lyon, France) with controlled food and water. At the end of this period, the food consumption was measured, and the 24h-calories intake was then calculated.

### Insulin and glucose tolerance tests

Insulin sensitivity was evaluated with an insulin tolerance test (ipITT). A single intraperitoneal (ip) injection (1 IU kg-1 of body weight) of human recombinant insulin (Umuline®, 100 UI ml-1, Lilly, Indianapolis, IN, USA) was delivered after a 4h-fast. Glucose tolerance was estimated with a glucose tolerance test (ipGTT) using a single ip injection of a D-glucose solution (100 mg/100 g of body weight) after an overnight fast. Blood glucose was then measured from the tail vein just before and 15, 30, 60 and 120 minutes after insulin or glucose injection, using test strips and reader (ACCU-CHEK Go®, Roche, Basle, Switzerland). The area under curves (AUC) were calculated using the trapezoidal rule of Prism 9 software (GraphPad Prism® Software).

### Plasma analysis

Total cholesterol, high-density lipoprotein (HDL), low-density lipoprotein (LDL), triglycerides (TG), alanine aminotransferase (ALT) and aspartate aminotransferase (AST) were measured using a C16000 chemistry analyzer (Abott®, Chicago, IL, USA) at the Biochemistry Laboratory of the University Hospital of Angers. Insulin was measured using the Rat Ultrasensitive Insulin ELISA kit (80-INSRT-E01, Eurobio, Courtaboeuf, France) according to the manufacturer’s instructions. Leptin was measured using an ELISA kit (mouse, rat) (#A05176.96, Bertin, Montigny-le-Bretonneux, France).

### Vascular reactivity of thoracic aorta and mesenteric arteries

Vascular reactivity was assessed by studying the reactivity of 4-months old rats isolated thoracic aorta and second-order mesenteric arteries. Four segments (2mm long) of thoracic aorta and mesenteric arteries from 1 animal were dissected and mounted on 2 wire-myograph (DMT, Aarhus, Denmark). Then 2 tungsten wires (25μm diameter) were inserted into the lumen of the arteries and fixed respectively to a force transducer and a micrometer. All arterial segments were bathed in a 5ml organ bath containing a physiological salt solution (130.0mM NaCl, 3.7mM KCl, 1.2mM MgSO_4_, 14.9mM NAHCO_3_, 1.6mM CaCl_2_, 5.0mM HEPES, 1.2mM KH_2_P0_4_ and 11mM glucose) maintained at a pH of 7.4, a pO2 of 160mmHg and a pCO2 of 37mmHg. After wall tension normalization at 90mmHg to approximate the *in vivo* transmural pressure[19,20], arterial segments were allowed to stabilize for 30 minutes. The ability of the 4 segments of thoracic aorta and the 4 segments of mesenteric arteries to contract was first tested with KCl (80mM) and secondly with cumulative concentration-response curve to phenylephrine (PE 1nM to 10µM). Then the integrity of the endothelium was tested using acetylcholine (ACh, 1µM) after precontraction with PE (1µM). After washout, the 4 arteries were precontracted with PE to a level approximately equivalent to 80% of the maximal response and vasorelaxation was assessed with a cumulative dose-response curve of ACh (1nM-10µM). Then, ACh-induced vasorelaxation was tested in the presence of *N* ^G^-nitro-L-arginine methyl ester (L-NAME, 100μM, nitric oxide synthase inhibitor), indomethacin (INDO, 10µM, nonselective inhibitor of cyclooxygenase enzymes COX1 and 2), KCl25 (25mM, EDHF blockade) and finally ACh-induced vasorelaxation was tested in presence of Tempol + Catalase (ROS inhibitor). At the end of the experiment, endothelium-independent vasorelaxation was studied with a cumulative dose-response curve of sodium nitroprusside (SNP, 1nM-10µM). All reagents were purchased from Sigma-Aldrich (Saint-Louis, MO, USA). Data were expressed as a % relaxation of PE-induced precontraction.

### Histological analysis

#### White adipose tissue

Visceral (perigonadal and perirenal) and subcutaneous (inguinal) white adipose tissues were collected; each fat pad mass was determined then fixed in 4% paraformaldehyde during 24h before embedded in paraffin. Adipocyte size was measured on 5µm-tick sections stained with hematoxylin-eosin (H&E) at HIFIH laboratory (Angers University). We used the Aperio CS2 digital slide scanner to generate a virtual slide with high quality images (resolution of 0.5 µm/pixel). For each animal, one slide was scanned and a region of interest (ROI) of 3500×3500 pixels was manually selected on the virtual slide. We took care to choose an artifact-free ROI to facilitate the automatic adipocytes detection. Then, adipocyte sizes were automatically measured in the selected ROI by our software (added as a Plugin in ImageJ).

The first step of the algorithm was to detect the adipocytes represented in high intensity (near white color) in H&E staining. Thus, we obtained a binary image (white and black) via an automatic thresholding technique developed in HIFIH laboratory. The other step consisted in separating aggregates of adipocytes using circularity criteria. Finally, we calculated the mean diameter of each adipocyte detected.

##### Liver

A sample of the median lobe of the liver was fixed in 4% paraformaldehyde during 24h before embedded in paraffin. Consecutive tissue 5µm-tick sections were realized and stained with 0.1% picrosirius red. Image acquisition was done with Aperio CS2 digital slide scanner (Leica Biosystem, Nanterre, France). Liver steatosis and fibrosis area were obtained by a semi-automatic detection using an algorithm developed by HIFIH laboratory (Angers). Liver steatosis correspond to the ratio of steatosis vacuoles/area of the whole biopsy sample [21].

##### Thoracic aorta and mesenteric artery

One segment of thoracic aorta and 1 second-order mesenteric artery were embedded in Tissue-Tek (Sakura Finetek, Alphen aan den Rijn, The Netherlands) and frozen in isopentane. Sections of 7µm thick were stained with orcein to measure histomorphometric parameters: external and internal media diameters, medial cross-sectional area (MCSA) and intima-media thickness (IMT), after image acquisition (Aperio CS2 digital slide scanner, Leica Biosystem, Nanterre, France) and analyze using ImageJ software (NIH, USA). Media-to-lumen ratio was calculated.

### Statistical analysis

All results are presented as means ± SEM compared using Mann-Whitney test for Control and HFHS comparisons, and a non-parametric test (Kruskal-Wallis) followed by Dunn’s post-hoc test for *Control*, *Obese* and *Sleeve* comparisons. Body weight and vascular reactivity curves were analyzed using a two-way ANOVA followed by a Bonferroni’s multiple comparisons test. Values of p<0.05 were significant. All statistical analyses were performed using Prism 9 software (GraphPad Prism® Software, San Diego, CA, USA), including comparison of variances.

## Results

### Impact of vertical sleeve gastrectomy on body weight and plasma metabolic profile of female rats 1 month after surgery

Minimum 8 weeks of HFHS diet led to a significant body weight gain of female rats compared to those fed with control diet (Fig. 1S-A). This was correlated with an increase of 24h-calories intake (Fig. 1S-B). The day of surgery, *Obese* and *Sleeve* female rats weighted 372.5 ± 8.8g and 361.4 ± 5.8g, compared to *Control* female rats (326.8 ± 6.5g, p<0.01). Sleeve gastrectomy induced a 10%-drop of body weight during the 5 days following surgery. Afterwards, *Sleeve* female rats maintained a similar body weight as *Control* female rats, although *Obese* female rats continued to gain weight (Fig. 1A). Interestingly, 24h-calories intake measurement showed a non-significant increase in *Obese* and *Sleeve* groups 1 month after surgery (Fig. 1B).

**Figure 1.**
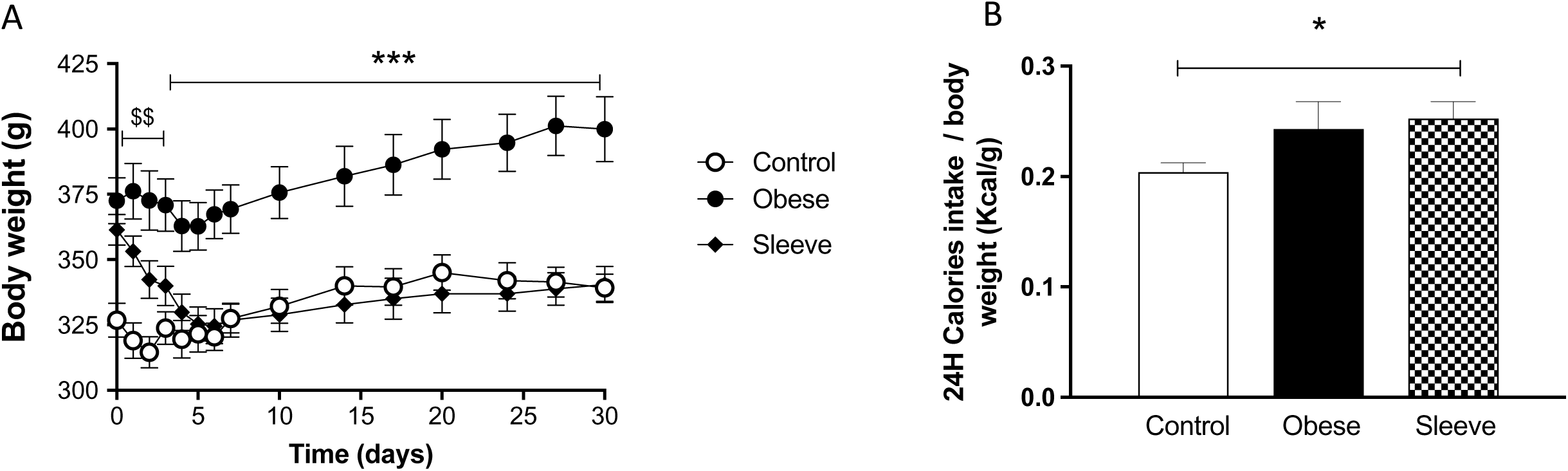
Impact of bariatric surgery on female rats 1 month after surgery. *(A)* Body weight follow-up during the period of 1 month after surgery, *(B)* Daily calories intake for *Control* (n=11), *Obese* (n=10) and *Sleeve* (n=13) female rats. Values are means ± SEM. ^$$^P<0.01 *Control* vs other mothers; *P<0.05 *Sleeve* vs *Control* mothers and ***P<0.001 *Obese* vs other mothers.

In addition, although total cholesterol, HDL and LDL were similar between groups, *Obese* female rats showed a glucose intolerance associated to decreased fast blood glucose, plasma TG levels and an increase of insulin and leptin levels. Sleeve gastrectomy decreased insulin plasma level and normalized leptin level compared to *Obese* female rats (Table 1).

**Table 1:** Impact of bariatric surgery on glucose metabolism and lipid profile of female rats 1 month after surgery. (*\* P*<0.05 and ** *P*<0.01 *Obese* or *Sleeve* groups vs *Control* group; ^$$^ *P*<0.01 *Sleeve* group vs *Obese* group. Data are mean ± SEM, n= 8 per group).

|  | Control | Obese | Sleeve |
| --- | --- | --- | --- |
| Blood glucose (mM) | 7.23 ± 0.53 | 5.40 ± 0.05* | 6.88 ± 0.46\$\$ |
| ipGTT |  |  |  |
| *Glucose AUC (0-120 min)* | 13466.2 ± 331.0 | 14940.2 ± 281.9** | 1417.5 ± 317.8** |
| Cholesterol (mM) | 2.38 ± 0.32 | 1.92 ± 0.12 | 1.83 ± 0.08 |
| *LDL Cholesterol (mM)* | 1.34 ± 0.22 | 1.19 ± 0.06 | 1.16 ± 0.04 |
| *HDL Cholesterol (mM)* | 0.67 ± 0.17 | 0.61 ± 0.15 | 0.54 ± 0.10 |
| Triglycerides (mM) | 0.80 ± 0.13 | 0.48 ± 0.07* | 0.43 ± 0.05* |
| Insulin (ng/mL) | 0.71 ± 0.19 | 1.54 ± 0.55 | 1.21 ± 0.39 |
| Leptin (ng/mL) | 1.44 ± 0.02 | 4.14 ± 0.14 | 1.28 ± 0.03 |

### Impact of vertical sleeve gastrectomy on body weight and plasma metabolic profile of female rats 4 months after surgery

Four months after surgery, *Sleeve* female rats retained a body weight like *Control*, although *Obese* female rats continued to gain weight (Fig. 2A). This result was not due to a growth perturbation because we did not detect difference in tibia length or tibial muscle weight between the 3 groups (Fig. 2A). In the *Obese* group, the increase of body weight was correlated to a subcutaneous / visceral tissues weight associated to adipocyte hypertrophy demonstrated by a shift in the size distribution curve of adipocytes (Fig. 2C-D). In the *Sleeve* group, even if female rats ate more (Fig. 2B), the normalization of body weight was associated with a decreased weight of white adipose tissue and a normalization of the size distribution curve of adipocytes compared to *Obese* female rats (Fig. 2C-D).

**Figure 2.**
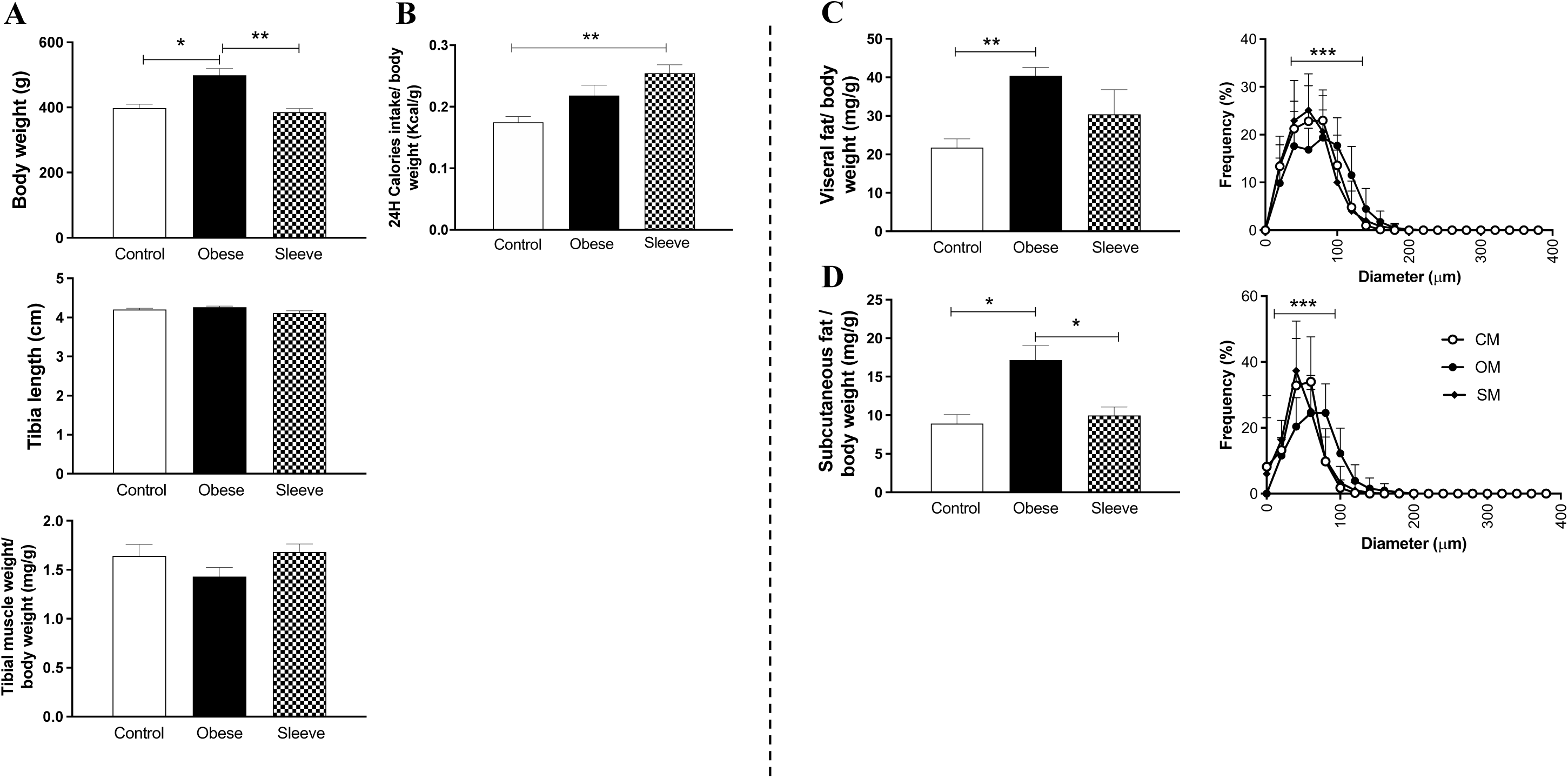
Impact of bariatric surgery on morphometric parameters and white adipose tissue of female rats 4 months after surgery. *(A)* Morphometric parameters with body weight, tibia length and tibial muscle weight, *(B)* Daily calories intake, *(C)* Visceral fat weight (left panel) and associated adipocyte size distribution curve (right panel) and *(D)* Subcutaneous fat weight (left panel) and associated adipocyte size distribution curve (right panel) in *Control* (n=6), *Obese* (n=8) and *Sleeve* (n=8) female rats. Values are means ± SEM. *P<0.05 and **P<0.01. ***P<0.001 *Obese* vs other mothers

At this time point, sleeve gastrectomy also normalized glucose tolerance, triglycerides, insulin and leptin levels compared to *Obese* female rats (Table 2).

**Table 2:** Impact of bariatric surgery on glucose metabolism and lipid profile of female rats 4 months after surgery. (* P<0.05 *Obese* vs *Contro*l group, ^$^ P<0.05 and ^$$^ P<0.01 *Sleeve* vs *Obese* group. Data are mean ± SEM, n= 7 per group).

|  | Control | Obese | Sleeve |
| --- | --- | --- | --- |
| Blood glucose (mM) | 6.23 ± 0.74 | 6.49 ± 0.71 | 5.23 ± 0.36 |
| ipGTT |  |  |  |
| *Glucose AUC (0-120 min)* | 13020.3 ± 354.6 | 15260.6 ± 334.1* | 14924.3 ± 409.2 |
| ipITT |  |  |  |
| *Glucose AUC (0-120 min)* | 5704.0 ± 575.2 | 6247.7 ± 433.3 | 5642.4 ± 157.7 |
| Cholesterol (mM) | 1.80 ± 0.14 | 1.56 ± 0.10 | 1.73 ± 0.22 |
| *LDL Cholesterol (mM)* | 0.88 ± 0.13 | 0.72 ± 0.08 | 1.14 ± 0.20 |
| *HDL Cholesterol (mM)* | 0.37 ± 0.07 | 0.31 ± 0.05 | 0.27 ± 0.03 |
| Triglycerides (mM) | 0.89 ± 0.06 | 1.44 ± 0.17 | 0.71 ± 0.17\$\$ |
| Insulin (ng/mL) | 0.77 ± 0.21 | 2.55 ± 0.83* | 0.78 ± 0.22\$ |
| Leptin (ng/mL) | 3.67 ± 0.04 | 10.79 ± 3.15 | 3.65 ± 0.26 |

### Impact of vertical sleeve gastrectomy on hepatic profile of female rats 4 months after surgery

Before surgery, *HFHS* female rats showed a higher percentage of steatosis area compared to *Control* female rats; fibrosis areas were not different between the 2 groups of female rats (Fig. 1S-C). Four months after surgery, although there is no difference in liver weight between groups (Fig. 3A), we still noted a weak percentage of steatosis in *Obese* group liver. Interestingly, we also detected liver fibrosis without steatosis in *Sleeve* group (Fig. 3B). This was associated with a significant elevation of AST in plasma compared to other groups (83.1 ± 7.3 U/L vs 59.7 ± 2.2 U/L for *Control* group and 79.6 ± 6.9 U/L for *Obese* group, p<0.05). In addition, while ALT was not detectable in plasma of *Control* group, we observed an increased serum level of ALT in *Obese* et *Sleeve* groups (14.2 ± 3.1U/L and 12.6 ± 0.9 U/L).

**Figure 3.**
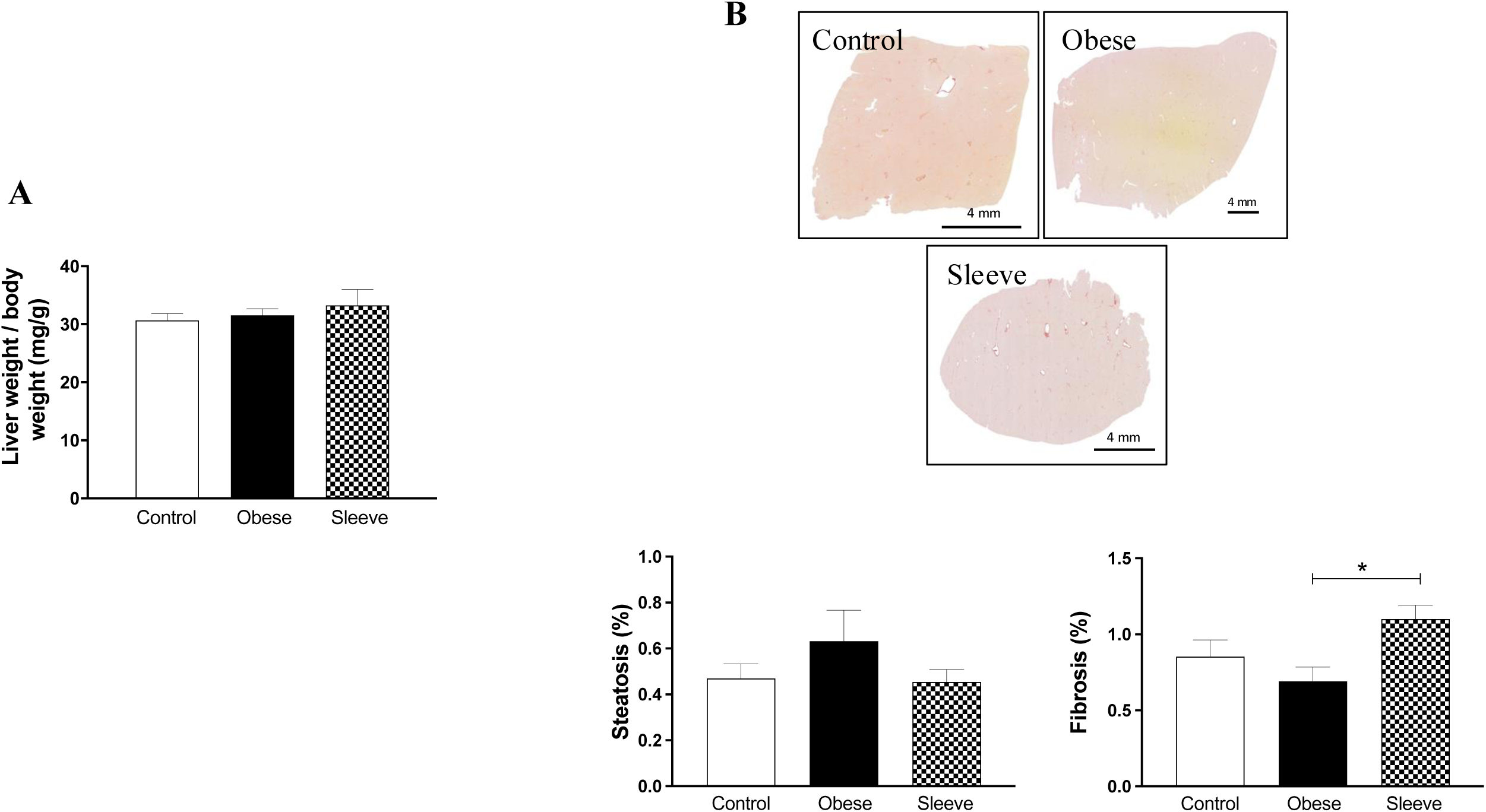
Impact of bariatric surgery on liver of female rats 4 months after surgery. *(A)* Rat liver weight, and *(B)* Examples of light microscopy images (scale bar = 4mm) and percentage of steatosis and fibrosis area of livers in *Control* (n=8), *Obese* (n=8) and *Sleeve* (n=7) female rats. Values are means ± SEM. *P<0.05 *Sleeve* vs *Obese* mothers.

### Impact of vertical sleeve gastrectomy on vascular profile of female rats 4 months after surgery

To evaluate the impact of bariatric surgery on the vascular profile, we studied vascular function (i.e. vascular reactivity) and structure of large (thoracic aorta) and small (mesenteric) arteries of female rats 4 months after surgery.

Vascular reactivity of thoracic aortas of *Control*, *Obese* and *Sleeve* female rats were not different. In fact, the smooth muscle cell-dependent vasocontraction induced by KCl or PE and the endothelium-dependent (Ach) or -independent (SNP) vasorelaxation were similar between the 3 groups of mothers (Fig. 4, left panel). Despite this absence of modification in vascular reactivity, we measured a significant increase of MCSA associated with a high Media-to-Lumen ratio of thoracic aorta in *Sleeve* group compared to the other groups (Table 3). These modifications of structure indicate the presence of a hypertrophic remodeling induced by SG.

**Figure 4.**
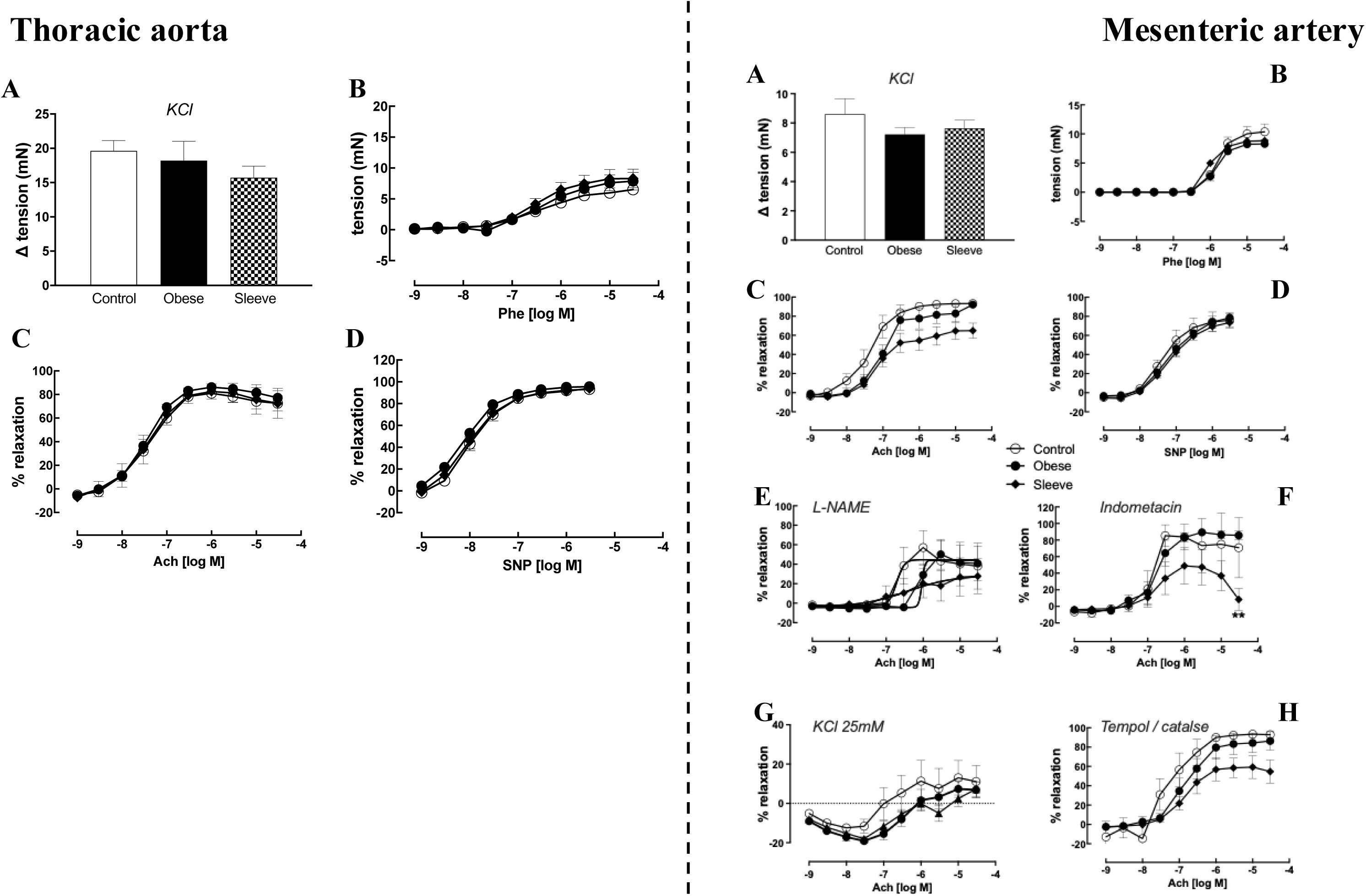
Impact of bariatric surgery on vascular function of female rats 4 months after surgery. Vascular reactivity of thoracic aorta (left panel) and mesenteric arteries (right panel) in *Control* (n=6), *Obese* (n=8) and *Sleeve* (n=6) female rats. Vasoconstriction response to *(A)* KCL 80mM and *(B)* cumulative doses of phenylephrine (Phe). Vasodilation responses to *(C)* cumulative doses of acetylcholine (ACh), *(D)* cumulative doses of sodium nitroprusside (SNP), cumulative doses of ACh in presence of *(E) N* ^G^-nitro-L-arginine methyl ester (L-NAME), *(F)* Indomethacin (INDO), *(G)* KCl 25mM (KCl25) and *(H)* Tempol + Catalase. Values are means ± SEM. Comparisons were made using two-way ANOVA with Bonferroni post-hoc test. **P<0.01 *Sleeve* vs *Obese* mothers.

**Table 3:** Histomorphometric analysis of thoracic aorta and mesenteric artery of female rats 4 months after surgery. Measurements of internal and external diameters, media cross-sectional area (MCSA), Intima-media thickness (IMT) and Media-to-Lumen ratio (*Control*: n=5, *Obese*: n= 6 and *Sleeve*: n=8; ** P<0.01 *Sleeve* vs *Control* group). Data are mean ± SEM.

|  | Control | Obese | Sleeve |
| --- | --- | --- | --- |
| ***Thoracic aorta*** |  |  |  |
| External diameter ($\mu\text{m}$) | 1799.5.2 ± 27.6 | 1889.2 ± 70.9 | 1931.9 ± 71.8 |
| Internal diameter ($\mu\text{m}$) | 1631.5 ± 32.6 | 1706.2 ± 90.9 | 1722.3 ± 77.1 |
| MCSA ($\mu\text{m}^2$) | 4532143.1 ± 17776.0 | 516246.4 ± 32525.0 | 597080.1 ± 25722.0** |
| IMT ($\mu\text{m}$) | 86.3 ± 3.4 | 97.3 ± 6.9 | 101.1 ± 5.0 |
| Media-to-lumen ratio | 0.217 ± 0.008 | 0.229 ± 0.017 | 0.265 ± 0.024 |
| ***Mesenteric artery*** |  |  |  |
| External diameter ($\mu\text{m}$) | 167.4 ± 17.5 | 189.2 ± 14.5 | 193.0 ± 11.5 |
| Internal diameter ($\mu\text{m}$) | 123.1 ± 13.8 | 141.3 ± 13.1 | 129.9 ± 4.3 |
| MCSA ($\mu\text{m}^2$) | 10919.6 ± 2817.3 | 10494.9 ± 1943.4 | 16619.7 ± 3672.5 |
| IMT ($\mu\text{m}$) | 23.2 ± 7.1 | 19.9 ± 1.9 | 26.8 ± 4.5 |
| Media-to-lumen ratio | 0.98 ± 0.24 | 1.13 ± 0.27 | 1.31 ± 0.32 |

In mesenteric artery segments, although KCl- or PE-induced vasocontraction was similar between the 3 groups (Fig. 4A and B, right panel), *Sleeve* female rats showed a lower response to Ach-induced vasorelaxation than *Control* group (Emax: 64.9 ± 7.9% vs 93.4 ± 2.7% for *Control* group, p=0.08, Fig. 4C, right panel) without alteration in endothelium-independent (SNP) vasorelaxation (Fig. 4D, right panel); this result highlights an endothelial dysfunction of mesenteric arteries of *Sleeve* female rats. In addition, blocking nitric oxide (NO) production by L-NAME induced a decrease of maximal response to ACh-induced vasorelaxation around 40% in *Control*, *Obese* and *Sleeve* female rats showing an equivalent implication of NO in mesenteric artery’s vasodilation for these 3 groups (Fig. 4E); nevertheless, we noted an increase of the EC50 in *Obese* female rats compared to *Control* group (1.094×10^-6^ ± 0.219×10^-6^ vs 0.127×10^-6^ ± 0.062×10^-6^, p<0.05), while this parameter was normalized in *Sleeve* female rats. The contribution of the prostaglandins (PGI_2_) pathway was then evaluated with INDO; although the vasodilator response to ACh in presence of INDO was similar between *Control* and *Obese* female rats, we observed an altered ACh-induced vasorelaxation with a reduced maximal response between *Sleeve* and *Obese* groups (-49%, *p<0.01*, Fig. 4F). This observation suggests a stronger involvement of the COX/PGI_2_ pathway in mesenteric artery of *Sleeve* female rats. The implication of the EDHFs pathway in endothelium-dependent vasorelaxation (explored with KCl 25mM) was not different between groups (Fig. 4G). Finally, in presence of Tempol + Catalase, Ach-induced vasorelaxation curves were equivalent to those without Tempol + Catalase (Fig. 4H) showing that ROS did not impact vascular reactivity of mesenteric arteries. In parallel, histomorphometric analysis of mesenteric arteries did not show significant modification even if all the parameters, except internal diameter, seemed to be increased for *Sleeve* female rats in favor of an outward hypertrophic remodeling (Table 3).

### Impact of vertical sleeve gastrectomy on offspring delivery and before weaning

During each birth, we measured the number of pups/litters. Even if the decrease was not significative, we observed a negative impact of maternal obesity on litter size, which is reversed by sleeve gastrectomy (Fig. 5A). This effect did not impact the sex ratio distribution or the birth weight of pups (Fig. 5B). But the body weight follow-up shows that OMO gained more weight until weaning than CMO and SMO males and females (Fig. 5C).

**Figure 5.**
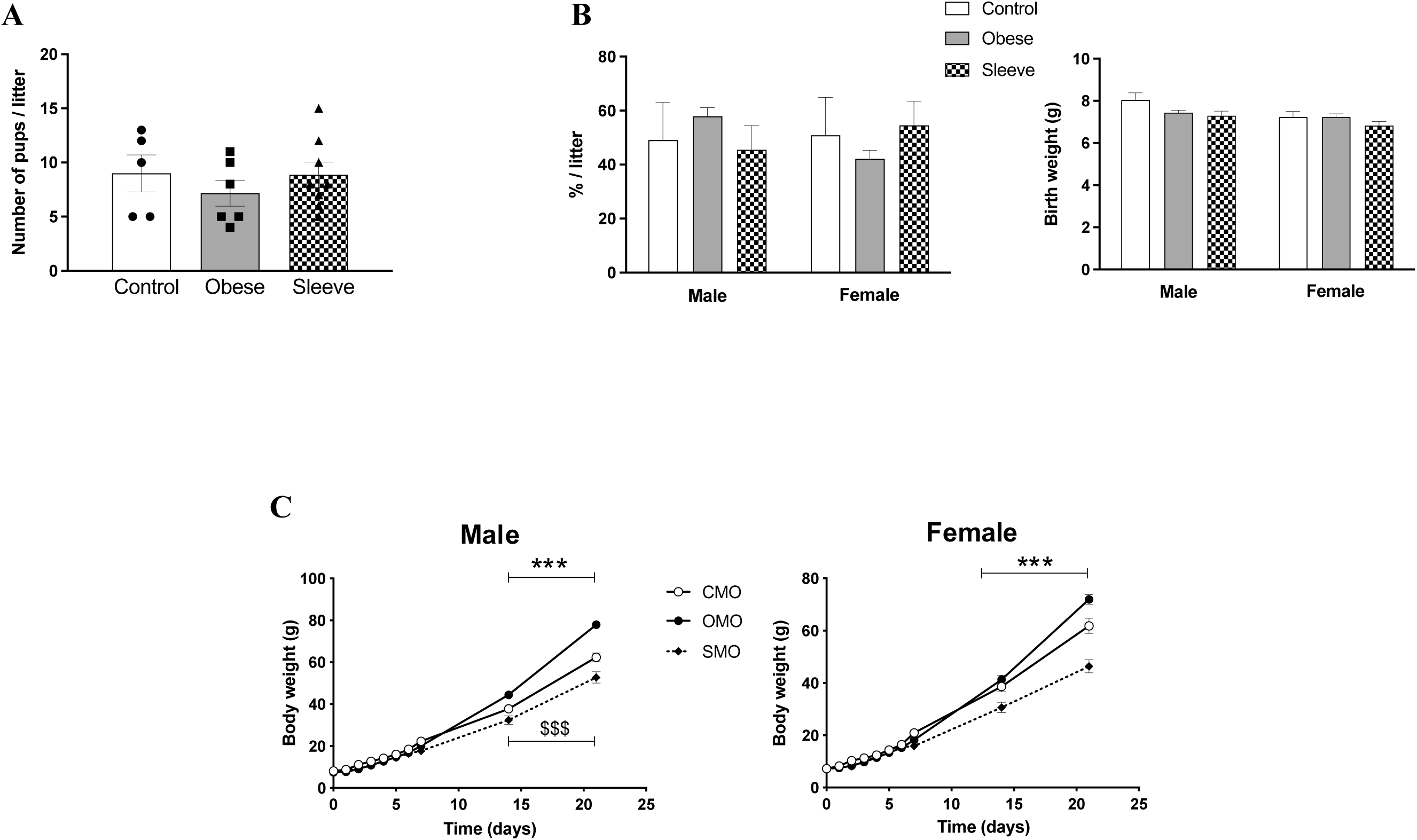
Impact of bariatric surgery on offspring before weaning. *(A)* Number of pups by litter and *(B)* Sex ratio distribution by litter for *Control* (n=5), *Obese* (n=7) and *Sleeve* (n=8) mothers. *(C)* Body weight follow-up during a period of 23 days before weaning of *Control mother offspring* (CMO, n=21), *Obese mother offspring* (OMO, n=23), and *Sleeve mother offspring* (SMO, n=16). ***P<0.001 OMO vs CMO and ^$$$^P<0.001 OMO vs CMO.

## Discussion

In less than twenty years, sleeve gastrectomy has become the most used technique for weight loss regarding its relative safety and efficiency [22]. Using an experimental surgical model on HFHS-induced obesity, we highlight here that female rats that have undergone SG show vascular alterations including endothelial dysfunction in mesenteric arteries and hypertrophic remodeling 4 months after surgery. In addition, even if these animals do not present liver steatosis unlike *Obese* female rats, they develop a higher level of fibrosis.

Our model of HFHS-induced obesity before surgery induces a 12% increase in weight gain associated with a 20% increase in caloric intake, resulting in a 3-fold increase of circulating leptin, impaired glucose tolerance and appearance of fatty liver and fat mass. It thus highlights the development of obesity and associated comorbidities in our animal model, even on a short time, which is not the case for most currently published rodent models of bariatric surgery [23,24]. In Human adults, guidelines currently recommend bariatric surgery when body mass index > 35kg/m^2^ [25,26] which approximately corresponds to a 40% body weight gain. This is more than the 12% increase of body weight in our model, but there is rising concern to propose this treatment even in patients with obesity and severe diabetes mellitus [27].

Nevertheless, our model mimics clinical effects of SG. Our preclinical study shows a rapid decrease of adipose tissue, restoration and maintenance of a normal body weight through gestation and lactation, despite a high food intake. Body composition analysis shows that this weight loss is due to a decrease in fat mass, more subcutaneous than visceral, without change in muscle mass as assessed by tibial muscle weight. Analysis of white adipose tissues shows the disappearance of hypertrophic adipocytes in both visceral and subcutaneous fat samples in the *Sleeve* group as compared to the *Obese* one. We also observe a normalization of leptin and insulin levels after one month with a sustained effect over time which may be partly explained by the endocrine function of adipose tissue [28]. As expected, postoperative weight loss was associated with improvement of GTT and ITT values and normalization of fasting insulinemia after 4 months.

In our study, we demonstrate that, 4 months after surgery, SM developed an altered endothelial function of resistance (mesenteric) arteries although thoracic aorta reactivity was preserved. The presence of an endothelial dysfunction after bariatric surgery is controversial; based on protein analysis of endothelin-1 axis on lung tissue, Ruze *et al*. recently concluded in endothelial function improvement after sleeve gastrectomy [29]. Conversely, it was also shown a decrease of plasma NO bioavailability 12 and 24 weeks after SG [30]. In our study, the altered vasodilator response to Ach seems not to involve NO and oxidative stress pathways since we did not detect any different effect with or without L-NAME or Tempol + catalase. But, the endothelial dysfunction of mesenteric arteries is linked to improved COX/PGI_2_ pathway, possibly prostacyclin. The surgical procedure involves removing a significant portion of the stomach. Then the alteration of endothelial function could have potential impact on oxygen delivery and nutrient absorption to the intestinal tissues potentially affecting their function and leading to gastrointestinal complications.

Oxygen is important for long-term control of local blood-flow. Interestingly we detected a vascular hypertrophic remodeling of both thoracic aorta and mesenteric artery in SM. Hypertrophic response increases the size of the vascular smooth muscle cells and stimulates additional extracellular matrix which is the hallmark of chronic hypertension later on. The detection of this structural modification was not expected since clinical data show more than 50% of hypertension remission one year after surgery [31]. Nonetheless, even if we did not measure blood pressure, heart and left ventricular weights were not different between groups (data unshown) pointing out an absence of elevated blood pressure. Moreover leptin has been described as a potential mediator of vascular remodeling associated with obesity [32], independently of blood pressure level [33]. Then, 4 months after SG, even if plasma leptin level was normalized, hypertrophic remodeling was not reversed. Thus our finding suggests a chronic increase of blood flow as observed in metabolic syndrome Zucker rats [34]. This hypertrophic remodeling could contribute to elevated blood flow in order to increase oxygen turn-over.

Concerning liver histology, sleeve gastrectomy was effective to normalize steatosis, but we observed a 2.5-fold increase in liver fibrosis as compared to *Obese* females, even if the level of fibrosis remained within the normal range. Bariatric surgery is known to improve MAFLD outcomes and fibrosis scores [35,36], but few studies have addressed this topic in the specific context of pregnancy and lactation. Pregnancy can cause diverse acute liver disorders [37] which usually resolve after delivery. In case of preexisting MAFLD or obesity, one recent study by Hussain *et al*. has found an increased liver stiffness after delivery with type 2 diabetes mellitus, preeclampsia and ALT ≥ 25 as risk factors [38]. In human, breastfeeding seems to be beneficial for liver prognosis, with a positive correlation between the length of breastfeeding and the lowering of the risk of developing MAFLD [39]; and on a longer follow-up, cumulative lifetime breastfeeding was related with a decreased likelihood of advanced liver fibrosis [40]. But recently, Zhang *et al.* have reported that during transition, dairy cows experienced an increase in liver fibrosis proportional to the underlying severity of liver steatosis [41]. Moreover, Bertasso *et al.* has found a worsening of MAFLD after Roux-en-Y gastric bypass in female rats after pregnancy and lactation [42]. Our results thus suggest the same liver impact after sleeve gastrectomy and need to be explored by further investigations.

Pre-conceptional bariatric surgery has proved it efficiency to reduce obesity-associated adverse obstetrical outcomes, but has also been associated with an increased risk of intrauterine grows restriction, small-for-gestational age infants and preterm deliveries [43,44]. Often reported after gastric bypass, similar results have been demonstrated with sleeve gastrectomy [45]. In our study, *Obese* mothers had a slightly lower number of pups/litter, may resulting from higher *in utero* deaths and/or from an altered fertility. On the contrary to previous finding on rodent offspring [46,47], our results show comparable birth weights of offspring born to *Sleeve* mothers than the ones born to *Obese* or *Control* mothers. Nevertheless, their subsequent growth curves before weaning showed an altered pattern in both males and females, in favor of an impaired nutritional intake, at least during lactation, whereas pups of *Obese* mothers grew faster.

Some limitations of our study should be discussed. Our model still has some divergent points from the clinical profile which could probably be due to only 8 weeks of HFHS diet before surgery. In fact this time period under HFHS diet is not sufficient to observe endothelial dysfunction in the *Obese* group as described in the literature [48]. But our aim was to generate female rats having undergone SG still of reproductive age at the period of mating. Then we chose as a marker of obesity development an elevation of body weight associated to a significant increase of plasma leptin level in *HFHS* group.

## Conclusion

The present study has highlighted the potential of sleeve gastrectomy performed on HFHS obesity-induced female rats to cause infraclinical dysfunction of the liver and disruption of vascular reactivity of the mesenteric bed. Regarding our results, our animal model may be sustainable to further explore the repercussions of sleeve gastrectomy on the mother-child couple long term follow-up.

## Acknowledgments

We thank Dr Khaled Messaoudi (Biochemical Department, University Hospital of Angers, France) who had performed plasma metabolic measurements, Cyril Le Corre and SCAHU (animal facility in Angers) to take care of the animals and Jean-Luc GrandPierre (HIFIH laboratory) who prepared all the liver samples before analyses.

---

**Figure. 1S.**
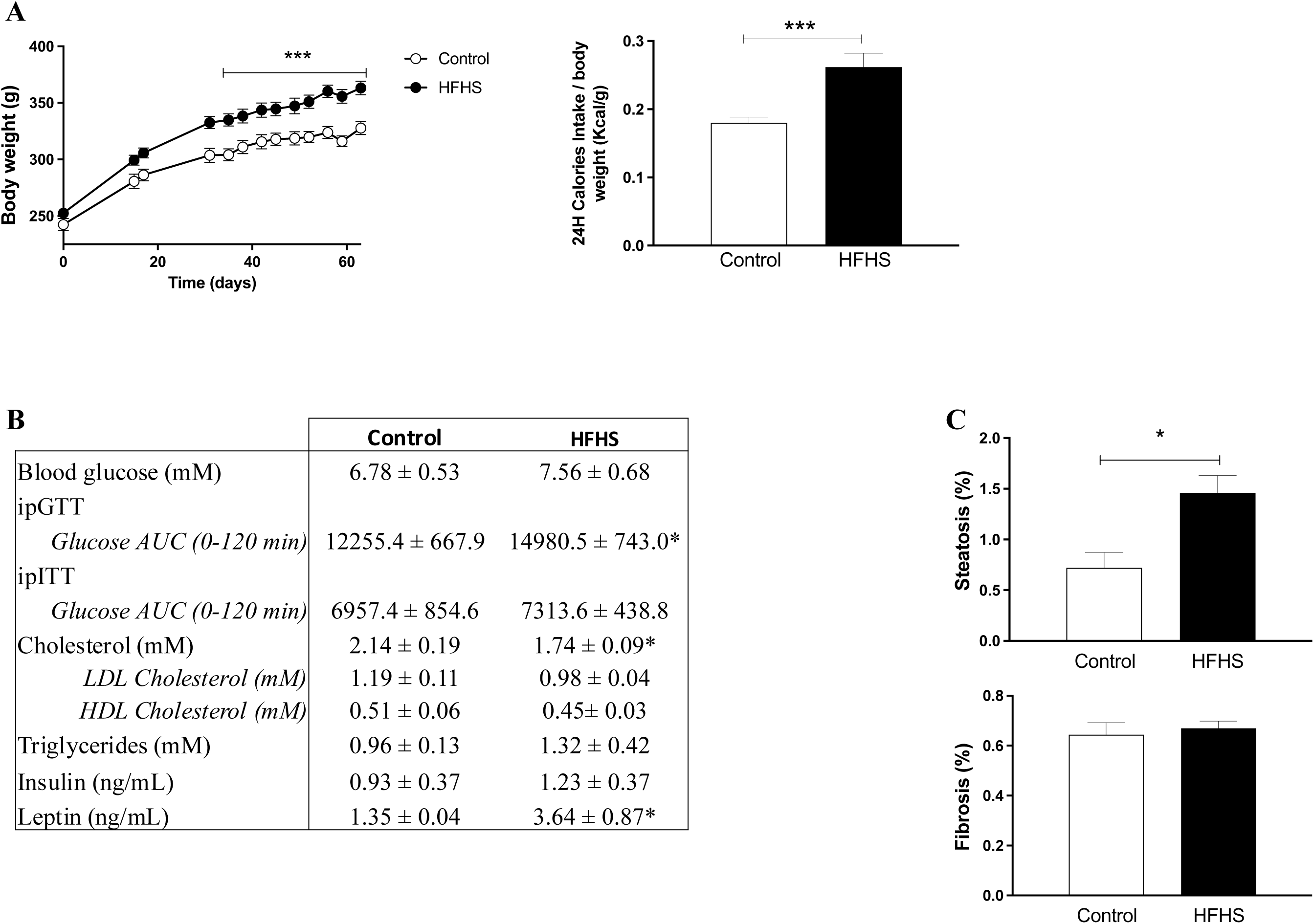
Experimental model of obesity on female rats. *(A)* Body weight follow-up from the beginning of high-fat / high-sugar (HF/HS) diet and daily calories intake after 8 weeks of diet for *Control* (n=11) and *HFHS* (n=24) female rats. *(B)* Glucose metabolism and lipid profile of *Control* and *HFHS* female rats (n=8 per group). *(C)* Percentage of steatosis and fibrosis area of livers on *Control* (n=11) and *HFHS* (n=24) female rats. Values are means ± SEM. *P<0.05 and ***P<0.001 *HFHS* vs *Control* female rats.

**Figure 2S.**
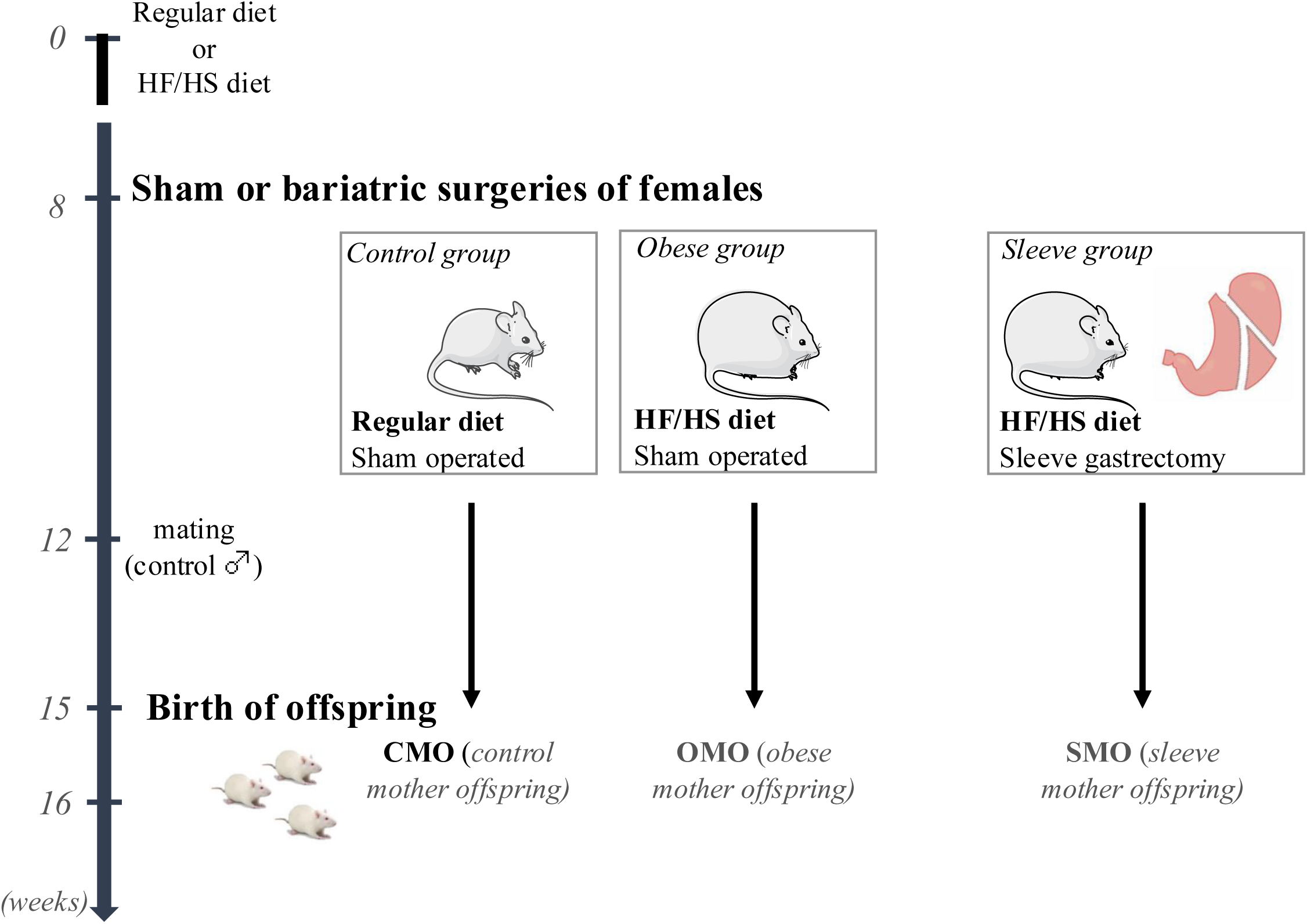
Experimental design of the study.

